# Information theory-based approach towards studying anti-coincidence detection via graded amplitude dendritic action potentials

**DOI:** 10.1101/2021.08.09.455769

**Authors:** Vidhi Sinha

## Abstract

In contrast to typical all or none action potential, recent discovery of graded amplitude action potentials in cortical neurons enabled the dendrites to perform XOR computation, previously thought to be performed only at network level. Thus, these special neurons can perform anti-coincidence detection at the dendritic level, but a lot is unanswered about this phenomenon. Can such experimentally observed dendritic action potential generating system transmit information about stimuli having varying degrees of temporal overlap? Can the system add to the repertoire of computations performed at dendritic level by enhancing the information transmission about varying amplitude stimuli? In this information theory-based study done in single compartment and two-compartment dendritic models, it is shown that such a system can indeed transmit information about the temporal overlap of stimuli as well as amplitudes of stimuli even at high input noise levels. First, the calculation of mutual information between single stimulus and response i.e. I(S;R) with varying noise showed that the information about temporally overlapping nature of stimuli is precisely transmitted by such a system. Secondly, the time evolution of mutual information was simulated through data from the system and it positively reinforced the above-mentioned result. Next, varying amplitude input stimuli was provided to the system and calculation of mutual information between two stimuli and one response i.e. I(S1,S2;R) with varying noise levels revealed that such a system optimally transmits the information about stimuli even at high noise levels. Finally, calculation of this information measurement with respect to time in an experiment with constant overlap but varying input amplitude again positively reinforced the result.

**Key Points:**

- Information theory-based measurements were employed to assess the role of graded amplitude dendritic action potentials.
- Action potentials (APs) with maximal amplitudes for threshold level stimuli and lower amplitudes for stronger stimuli were modelled with high voltage Ca2+ (HVA like) channels, Ca2+ dependent (BK-like) channel, leak channel and calcium pump in a single compartment model and two compartment dendritic model.
- Analysis done here, on comparison with control compartment generating constant amplitude AP via standard Hodgkin-Huxley sodium potassium channel revealed that such a compartment shows optimal information transmission about both varying amplitudes input current stimuli as well as varying time overlap stimuli.

## Introduction

Since the discovery of active conductance gradients across the dendritic tree, several possible computational functions have been suggested that can be performed by these active dendrites[1]–[3]. This includes implementation of logic gates[4]; learning mechanism for long-term potentiation[5]; affecting learning capacity of neuron[6] and performing coincidence or anti-coincidence detection[7]. Coincidence detection is the phenomenon by which a single neuron or neuronal circuit transmits information about temporally close inputs from spatially distributed pre-synaptic neurons[8]. Whereas anti-coincidence detection is the phenomenon by which temporal information is encoded by the neuron or neuronal circuit computing the anti-coincident function with pre-synaptic neuronal responses as inputs[9]. Thus, if the inputs are temporally close enough, the post-synaptic neuron suppresses its response i.e., it relies on the suppression of neuronal response when different presynaptic inputs are separated by less temporal distance.

The active properties of dendrites give them an immense repertoire of computational transformations done on synaptic inputs by them even in the case of all or none constant amplitude dendritic action potential[10][11]. But the discovery of special graded calcium-based dendritic action potentials (dCaAPs) which have the property of showing maximum response to threshold stimuli and then a decrease in response for increasing stimuli above the threshold stimuli, has enhanced this repertoire even more[9]. These dCaAPs were found in human layer 2-3 pyramidal neurons and it was revealed that such a system can perform biological gate XOR computation at dendritic level[9], a transformation earlier thought to be only possible at network level[12]. Since XOR computation essentially enables the system to perform ‘anti-coincidence’, it will be interesting to computationally model these APs using various active ion-channel conductances and see what kind of advantage it offers over a conventional system generating all or none APs[9]. Active properties of dendrites give them the ability to perform ‘coincidence detection’ of afferent inputs often seen in pyramidal neurons and this property has been extensively studied across systems in terms of its possible functions as well as the mechanisms, both experimentally and computationally[3]. Given the fact that this property proved to be so essential in sensory perception[13]–[16], temporal coding in auditory cortex[17], coding of amplitude modulated stimuli[18], plasticity[19] and a lot of other physiological roles[3], [20], it is reasonable to investigate what kind of functions its counterpart ‘anti-coincidence detection’ might play.

When the synaptic current inputs are given to a post-synaptic neuron, the information about the stimuli can be transmitted by variation in amplitudes of multiple inputs or by having different temporal distances across those inputs. Thus, this study helps in throwing light about the role of such a system generating dCaAPs in transmitting information about stimuli, both in terms of varying temporal and varying amplitude trials. Computational simulations were done on a simple single compartment model and a more realistic dendritic model for achieving that goal. To assess how well variability in temporal overlap and amplitude of inputs is transmitted by a neuron showing anti-coincidence phenomenon, Shannon’s mutual information measures were calculated[4], [21]. By calculation of mutual information between stimulus and response i.e. I(S,R) in the first set of computational simulations, it is shown that the system can have enhanced ability to precisely transmit information about stimuli having different degrees of temporal overlap as compared to control system generating constant amplitude APs. In the next set of experimental simulations, calculating mutual information between two stimuli and response i.e. I(S1,S2;R) for pairs of different amplitude stimuli, it was seen that the system can transmit information about such stimuli as compared to control even when the noise ratio to mean amplitude is very high. Finally, time evolution of these mutual information measures in both the set of experiments gives more insight about the information transmission across the simulation duration in the context of graded action potentials for the stimuli.

## Methods

The goal of this study is to understand the phenomenon of anti-coincidence detection in a simple system where such graded dendritic action potentials are generated with respect to information transmission. Thus, the model chosen is a single compartmental model inserted with various ion channels to simulate dCaAPs as mentioned in previous studies[9].Another two-compartment dendritic model has also been investigated to simulate the graded dendritic action potentials, to see how such a model’s architecture affects the phenomenon and information processing. Then a series of computational simulation-based experiments were done on both these models to better understand the role of this phenomenon in transmitting information regarding the temporal structure as well as the amplitude of multiple stimuli.

### Single Compartment Model

Single compartmental simulations were done in the NEURON simulation environment[22]. The compartment was of length 100μm and diameter 99 μm. The parameters and channels were chosen from supplementary information of the study by Gidon et al[9].It was inserted with high voltage Ca2+ (HVA-like) channels, Ca2+ dependent (BK-like), L-type Ca2+ channel, fast K+ channel and leak channel. The cm was chosen to be 0.45uF/cm2 and the rm was 14,000/cm i.e. 31.111kOhm-cm2. With the mentioned parameter values, Rin turned out to be 100MOhm (Rin = Rm/πdl), which matches the experimental value (Fig 1a). The model mentioned in the study had modified time constant for BK-like channel so that the experimentally observed kinetics of dCaAPs are matched. Similarly, the fast K+ channel has modified equations and additionally L-type Ca2+ channel was added to ensure that the first AP doesn’t have unreasonably high amplitude as mentioned in the supplementary information. Other parameters of the channel were again taken from the same study and the publicly available model for this study posted on http://modeldb.yale.edu[9[9]. The compartment replicated the experimentally observed phenomenon of graded dendritic action potential which has max amplitude for threshold stimuli and decreasing amplitude for increasing current injection values in contrast to typical all or none action potential or the conventional action potential that increases in amplitude with stronger stimuli (Fig 1b). The phenomenon of anti-coincidence can be clearly seen in the experimental compartment of the model. Here two current inputs were given with a fixed temporal overlap to visualise the anti-coincidence phenomenon. Since, the stimulus duration is 850ms, delay is 100ms and overlaps is 300ms, we can see that from 650ms the spike height has greatly reduced and all that can be seen is noise fluctuations (Fig 1c).

**Fig 1:**
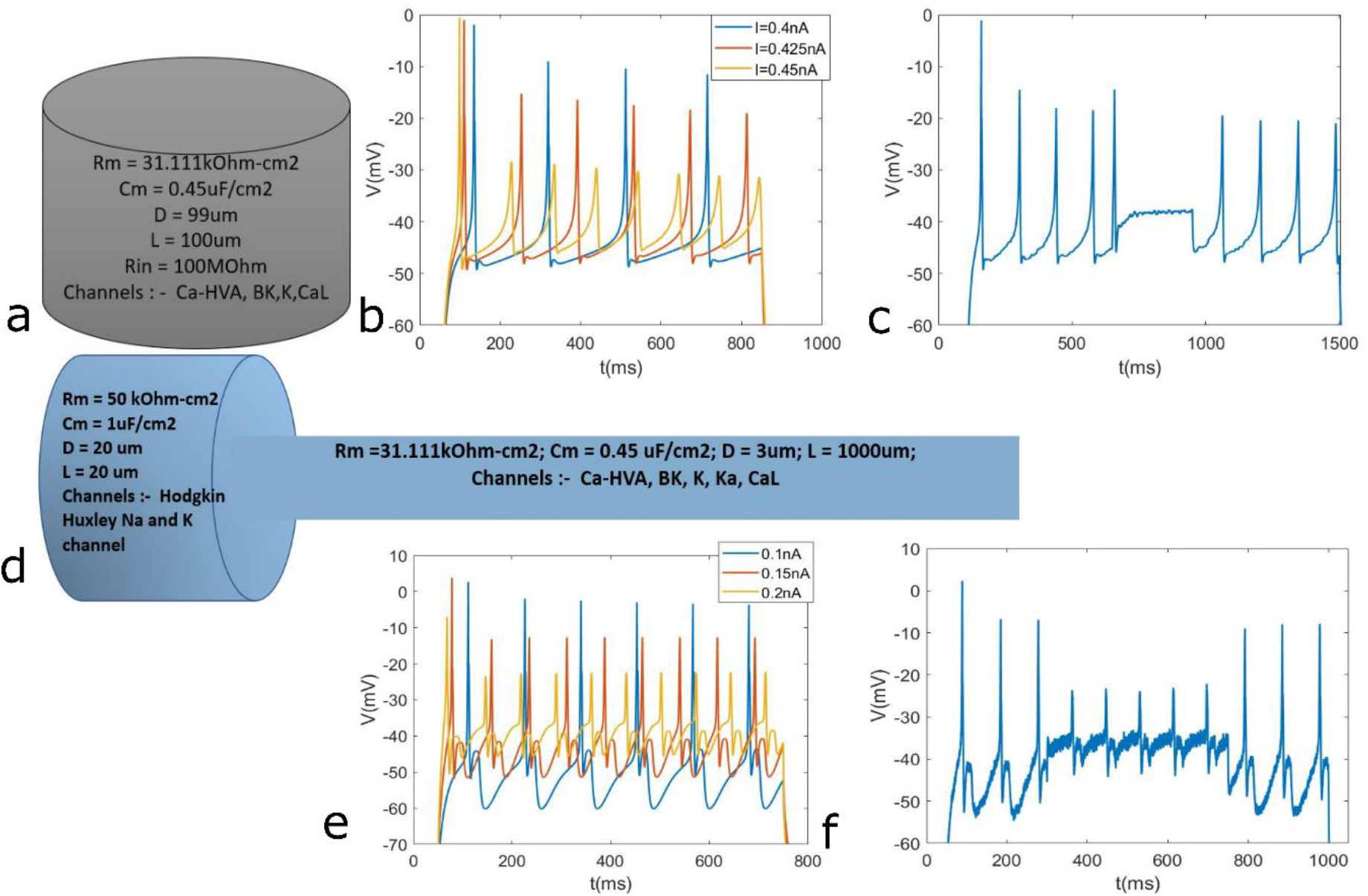
Model description: Panels (a) and (d) show the schematic of both models. Panels (b) and (e) show the working of graded AP model by showing response to increasing current amplitudes. Panels (c) and (f) show the response to two input current injected at different time points with a fixed temporal overlap to show the anti-coincidence. (a) Single compartment cylindrical model 99um diameter and 100um length with Rm of 31.111 kOhm-cm2 and Rin of 100MOhm. (b) Voltage response of the model to different input current. As the input current amplitude increases, the amplitude of the dCaAP decreases. (c) Phenomenon of anticoincidence shown for an overlap of 300ms, stimulus duration for both inputs as 850ms and delay for first input as 100ms where the noise value N was 0.2. (d) Two compartment dendritic model. Soma has 20um length and diameter, Rm of 50kOhm-cm2 and dendrite has length 1000um, diameter 3um and Rm of 31.111kOhm-cm2. (e) Voltage response of the dendritic model to different input current. As the input current amplitude increases, the amplitude of the dCaAP decreases. (f) Phenomenon where AP amplitude is greatly reduced, for an overlap of 450ms where stimulus duration for both inputs was 700ms and delay for first input was 50ms. The noise value N was 0.2.

Two types of control compartments were chosen. Both were inserted with Hodgkin-Huxley sodium and potassium channels. Control 1 has lower average firing rate i.e. less excitability and control 2 has higher average firing rate i.e. more excitability (Control 1, Average firing rate is 15 spikes/s with SD = 1 spike/s, gNaBar = 0.1 S/cm2, gKBar = 0.03 S/cm2; Control 2, Average firing rate is 55 spikes/s with SD = 2 spike/s, gNaBar = 0.25 S/cm2, gKBar = 0.05 S/cm2. The values of spike numbers are rounded off to integers. Here the mean current amplitude used was 0.425nA and the coefficient of variation (N) i.e. mean/SD for gaussian noise ranged from 0.2-3 as described later in experiments).

### Dendritic Model

Dendritic model simulations were also done in the NEURON simulation environment[22]. Soma was of length 20um and diameter 20um, whereas the dendrite was of length 1000um and diameter 3um. The soma was inserted with Hodgkin Huxley sodium and potassium channels. The dendrite was inserted with high voltage Ca2+ (HVA-like) channels, Ca2+ dependent (BK-like), L-type Ca2+ channel, fast K+ channel, KA channel and leak channel[23], [24]. The Ca-HVA, BK and CaL channels followed a step function distribution i.e. their conductances were uniform beyond 600um but for value of length lesser than 600μm, the conductances were zero. On the other hand, KA and fast K channels, followed a uniform distribution throughout the length of the dendrite. Rm of soma was 50kOhm-cm2 and Rm of dendrite was 31.111kOm-cm2. Maximum conductance values chosen for different channels are shown below-

**Table 1:** Channel Parameter Values.

| Sr No. | Channels | Maximal Conductance |
| --- | --- | --- |
| 1. | High Voltage Ca<sup>2+</sup> (HVA-like) channel | 0.0010 S/cm<sup>2</sup> |
| 2. | Ca<sup>2+</sup> dependent BK channel | 500 pS/μm<sup>2</sup> |
| 3. | L-type Ca<sup>2+</sup> channel | 0.05 S/cm<sup>2</sup> |
| 4. | KA channel | 0.00015 S/cm<sup>2</sup> |
| 5. | Fast K<sup>+</sup> channel | 0.3 S/cm<sup>2</sup> |

This dendritic compartment also showed the phenomenon of graded dendritic action potentials i.e. the amplitude of which decreased as the input current amplitude increased (Fig 1e). A less pronounced version (compared to single compartment model) of anti-coincidence phenomenon is also seen in this model. The stimulus duration is 700 ms, delay is 50ms and overlaps is 450ms, we can see that from 300ms the spike height has reduced a lot and we see noise fluctuations along with below threshold voltage depolarisations (Fig 1f). In this study, the dendritic model has also been used to shed further light on the effects of anti-coincidence phenomenon on information transmission.

Two control compartments were chosen for this model as well, where both of them had simple Hodgkin Huxley channels inserted. Soma channel conductances were kept identical in both the controls, but dendritic channels were varied. Control 1 had a lower average firing rate i.e. less excitability and control 2 had a higher average firing rate i.e. more excitability (Control 1, Average firing rate is 9 spikes/s with SD = 1 spike/s, gNaBar = 0.8 S/cm2, gKBar = 0.04 S/cm2; Control 2, Average firing rate is 40 spikes/s with SD = 1 spike/s, gNaBar = 0.435 S/cm2, gKBar = 0.02 S/cm2. The values of spike numbers are rounded off to integers. Here the mean current amplitude used was 0.425nA and the coefficient of variation (N) i.e. mean/SD for gaussian noise ranged from 0.2-3 as described later in experiments).

### Inter-pulse interval (IPI) overlap experiment

In this set of simulations, two noisy input currents were injected at 0.5 normalised location of the single compartment (1 normalised location on dendrite for dendritic model). For all the trials of this experiment the mean amplitude of the current was fixed at 0.4nA (0.15nA for dendritic model). The total simulation was run for 1800ms with dt = 0.025 i.e. 40 steps per ms. The delay to first current input was set as 100ms. Then the total duration for the input was set to 800ms. The total duration for the second current input was also 800ms but the delay for this input was set as-

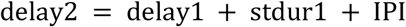

Where ‘delay1’ and ‘delay2’ are delays for first and second current input respectively; ‘stdur1’ is the stimulus duration for first current and ‘IPI’ is the inter-pulse interval. Here, the stdur1 was 800ms, fixed for all trials but IPI that denotes the amount of overlap between the two inputs, varied with every trial from −200ms to −1ms. Thus, each experiment consisted of 200 trials of uniformly distributed stimulus S which is mainly characterised by the IPI values. For one whole experiment, the coefficient of variation or noise ratio (N) was kept constant (N = SD/Stamp, where stamp is the mean current amplitude and SD is the standard deviation of input white noise current)

### Mutual Information calculation for IPI overlap experiment

Analysis was done on MATLAB after data collection from simulations. The joint probability distribution was created between stimulus S and response R. Here response R was considered to be the voltage. So, for a given experiment, voltage response values were binned in 1mV bins ranging from minimum to maximum of collected data for the whole experiment. S distribution was uniform with each stimulus being characterised by the overlap or IPI value, which was presented once. The formulae used for analysis to get the information theory measures are[21]-

1. H(R)= –∑p(r)log (p(r))
2. H(S)= –∑p(s)log (p(s))
3. H(R, S)= –∑p(r,s)log (p(r,s))
4. H(R|S)= H(R,S)– H(S)
5. I(R; S)= H(R)– H(R|S)

Each experiment was run 16 times with varying noise ratio N values ranging from 0 to 3.0 with spacing of 0.2, such that 16 different values of I(R;S) was calculated in both the control as well as experimental compartments, to get the parameter values for varying noise in the single compartment model. For the dendritic model, N values ranged from 0 to 1 with spacing of 0.1, such that 11 different values of I(R;S) was calculated for controls and experimental compartments. For getting the time evolving plot, 10ms resolution was taken and after every 10ms the voltage response values for next 200ms were collated together to calculate the I(R;S) for each of these points in both the control and experiment compartments for both the models. The plot was generated for constant noise ratio N for one set of simulations.

### Sensitivity Analysis

To make sure that the results were significant, sensitivity analysis was done on the parameters of the simulation. Mean current amplitude was varied form 0.375nA to 0.55nA for a wide number of values and representative noise varying and time evolving plots were shown for 0.4nA, 0.425nA and 0.45nA in the single compartment model. For the dendritic model, mean current amplitude was varied from 0.1nA to 0.2nA and representative noise varying as well as time evolving plots are shown for 0.15nA, 0.175nA and 0.2nA. For all the channels in both the models, the maximum conductance values were also varied. It was made sure that each parameter value was varied across a range covering at least 0.5 times to 2 times of original experimental compartment values.

### Varying amplitude stimuli experiment

Experiments in this section are done to study the information transmission in a system with stimuli that arrive for same duration but have different amplitudes. Here for the noise varying parameter calculation, both the stimuli were provided with a delay of 50ms. The duration of both the inputs S1 and S2 were fixed for 900ms, the simulation ran for 1000ms.

For single compartment model, S1 consisted of mean current amplitude varying from 0.1nA to 0.55nA in steps of 0.05nA. Same for S2, thus creating the (S1,S2) combination of (0.1,0.1),(0.1,0.15),(0.1,0.2)..(0.55,0.55). This ensured that S1 and S2 were individually uniform as well as making the (S1,S2) joint distribution to be uniform. For each set of experiment the noise ratio N was kept to be same.

For dendritic model, S1 consisted of mean current amplitude varying from 0.1nA to 0.4nA in steps of 0.025nA. S2 was also same, which made the (S1,S2) combination of (0.1,0.1),(0.1,0.125),(0.1,0.15)..(0.4,0.4). Thus, stimulus domain consisted of this S1,S2 joint distribution which mainly varied in their amplitudes. Here as well, each experiment had a fixed noise ratio N.

For getting the time evolving plot the experiment was adjusted a bit for both models, delay for the first stimulus was taken to be 50ms. Delay for the second stimulus was fixed at 300ms. Both stimulus S1 and S2 duration was for 700ms, thus keeping a fixed overlap of 450ms. The total simulation was run for 1200ms.

### Information theory measure calculation for Varying amplitude stimuli input experiment

A joint distribution of S1,S2,R was created where S1,S2 constituted the stimulus distribution from above and R was the voltage response. The voltage response R from all the possible stimuli conditions was pooled to get minimum and maximum voltage response values. For each stimulus condition i.e., S1 = k1, S2 = k2 (k1,k2 are possible stimuli values), R was binned from minimum to maximum voltage value with binwidth of 1mV. After getting the 3 variable joint distribution, marginalisation of the distribution was done to get various 1 variable and 2 variable joint distributions as needed.

Then following formula was used to calculate conditional mutual information I(S2;R|S1) [21]-

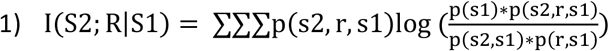

This was finally used to calculate I(S1,S2;R) with equation-

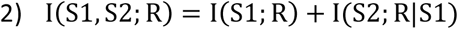

where I(S1;R) was calculated in the same as previously described.

For getting the time evolving plot, 5ms resolution was taken and after every 5ms the voltage response values for next 200ms were collated together to calculate the I(S1,S2;R) for each of these points in both the control and experiment compartment in the set of experiments where constant overlap of 450ms was considered. The plot was generated for constant noise ratio N for one set of simulations.

### Sensitivity Analysis

For all the channels in both models, the maximum conductance values were varied similarly here as well. It was made sure that each parameter value was varied in a range covering at least 0.5 times to 2 times of experimental compartment values. Then plots for varying noise and time evolution were made for representative values, same as that mentioned before. Temporal overlap window was also varied to make sure that the window itself doesn’t affect the conclusions.

### Statistical Test

For both the experiments, the noise mutual information values were collated and compared with control set after running the whole set of simulation twice, i.e getting 32 data points for both control and experiment in single compartment model and 22 data points for both control and experiment in dendritic model. Then unpaired t-test was performed on both the sets, where the test rejects the null hypothesis at 5%significance value, and the corresponding p-values were calculated using MATLAB.

## Results

### Dendritic calcium-based action potential generation in model

The dendritic calcium-based APs (dCaAPs) are generated because of the activation delay of about 2ms between the high voltage activated calcium channel and calcium dependent BK channel in the model, which can be a possible mechanism from the plethora of diverse possibilities, since these channels are known to express in human neurons. During the overlap period of anticoincidence phenomenon in this model, the conductance of calcium channel increases rapidly due to which calcium concentration increases. This affects the conductance of Ca-dependent BK channel which also increases in concentration that leads to high hyperpolarising current and thus preventing further AP generation. Such a model can then be used to study the effects of this phenomenon on information transmission, as done in this study.

### Inter-pulse interval overlap experiment

It is seen that as the noise fraction N increases, the mutual information between stimulus and response i.e. I(R;S) increases in both control as well experimental compartment, though that of experimental compartment is constantly higher than that of both higher excitable and lower excitable controls. On collating the calculated mutual information points of the whole duration of simulation for control and experimental compartments, the I(R;S) comes out to be significantly higher (p = 6.8651×10^−21 for control1 and p = 4.7088 × 10^−23 for control 2 in single compartment; p = 2.769×10^−11 for control1 and p = 2.4379 × 10^−13) in experimental compartment than that of control compartments in both the models (Fig 2a and Fig 2c). This just tells us that there is a large reduction in uncertainty about stimulus having different degrees of temporal overlap after observing the voltage response for the experimental compartment.

**Fig 2:**
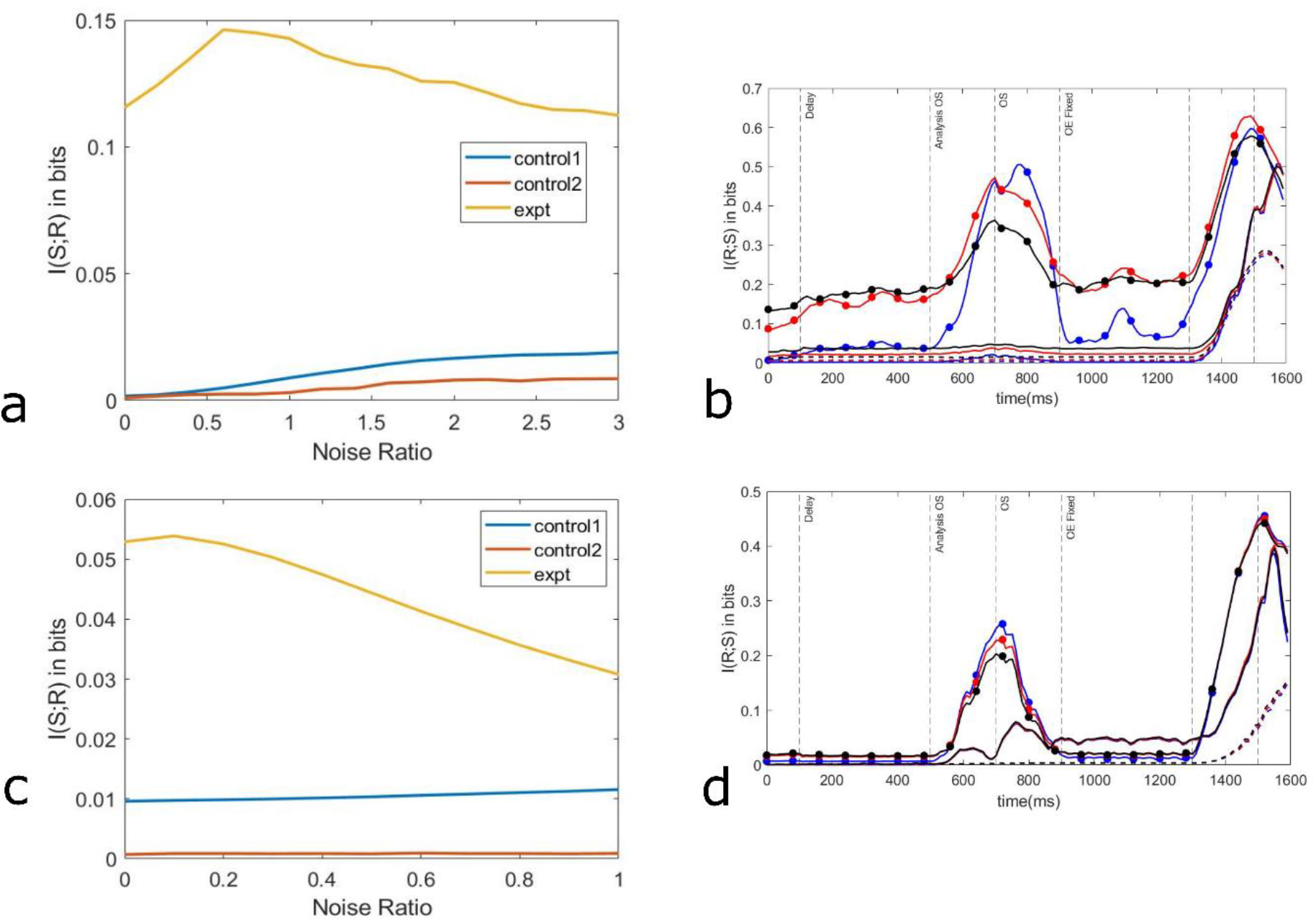
Results from IPI overlap experiment-Mutual information I(R;S) is plotted by collating the points from the restricted time of the simulation during which overlap occurred, with varying noise for experimental compartment and both the controls, results for single compartment model (a) and dendritic model (c) are shown. Time evolution of mutual information I(R;S) across the whole duration of the simulation for 3 different N values in single compartment model and dendritic model. ‘Colour’ represents the noise value N and ‘line style’ of each graph represents the nature of compartment i.e. experimental, control 1 or control 2. ‘Blue’ represents N=0.2 for single compartment model (b) and N=0.1 for dendritic model (d), ‘Red’ represents N=1 for single compartment (b) and N = 0.3 for dendritic model (d) and ‘Black’ represents N=2 for single compartment model (b) and N=0.5 for dendritic model (d). Graphs with ‘filled circle’ markers represent experimental compartment in each colour, those with ‘solid lines’ represent control 1 and those with ‘dashed lines’ represent control 2 for each unique colour i.e. different N value.

It can be seen from the graphs that I(R;S) for all noise values start increasing from 500ms and reach their stable lower value at 900ms (Fig 2b and Fig 2d). Even though the earliest overlap starts at 700ms, we see its effect at 500ms in the graph due to the nature of analysis done. Since response from 200ms later are already collated to generate the response, we see the increasing effect earlier from 500ms. On the other hand, the fixed value of overlap end at 900ms for stimuli make sure that I(R;S) stabilised to the lower value after increasing at the same time point i.e. 900ms (Fig 2b and Fig 2d). The increase in I(R;S) for the graphs make sense since during this particular region of increase, the stimuli takes their effect i.e. varying degrees of temporal overlap are observed. During the end part of graphs in terms of time, the I(R;S) values for both controls and experimental compartment rises, which is just because of variability in end time of second input current and the point at which the effect is seen in the analysis. When comparing the graphs for different N values to respective control compartments, it is observed that the mutual information time evolution peak increases during the region where stimuli temporal overlap’s effect is seen in analysis, as N decreases.

On varying the current amplitude for the IPI overlap experiment, it was observed from the noise varying and time evolving plots that as the current amplitude increased, the mutual information between stimuli and response decreased (Fig 3a and 3d for single compartment; Fig 4a and 4d for dendritic model). This is because of the nature of dCaAPs that show the anti-coincidence phenomenon, i.e., their amplitude decreases as the input current’s magnitude increases. This ultimately makes the information transmission about varying degrees of temporal overlap difficult when the current amplitude itself increases.

**Fig 3:**
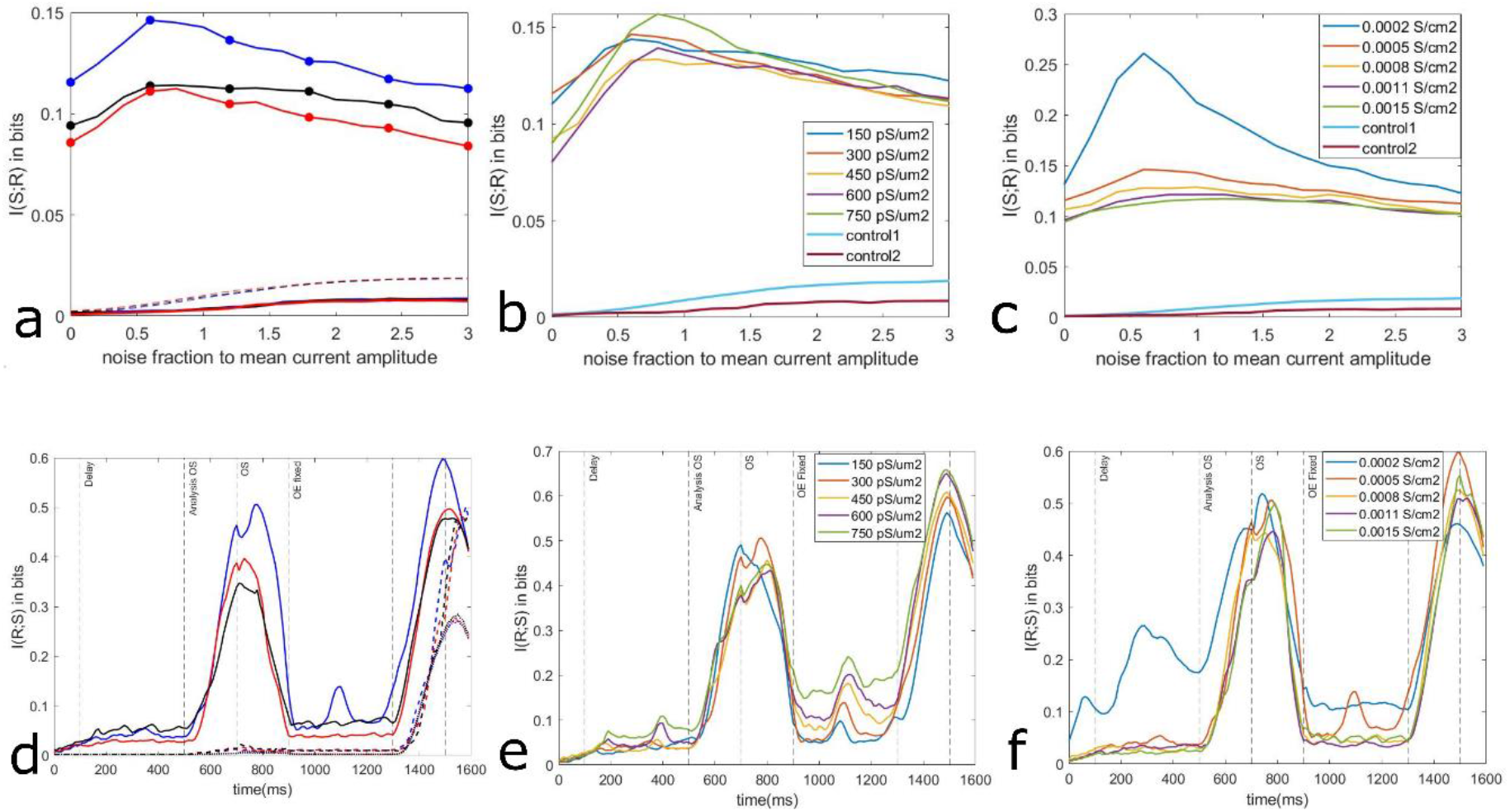
Sensitivity analysis for IPI overlap experiment in single compartment model a) Mutual Information I(R;S) is plotted for restricted time of simulation where overlap occurred with varying noise values, for different current amplitudes of the inputs. ‘Colour’ represents the current amplitude and ‘line style’ represents the nature of compartment i.e. experimental, control 1 or control 2. ‘Blue’ represents current amplitude of 0.4nA, ‘Black’ represents current amplitude of 0.425nA, ‘Red’ represents current amplitude of 0.45nA. Graph with ‘filled circle’ markers represent experimental compartment in each colour, ‘dashed line’ represents control 1 and ‘solid line’ represents control 2. Due to less variation in control compartments’ voltage response for different input current amplitudes, the differently coloured graphs are overlapping for both control compartments. b) Mutual Information I(R;S) is plotted for restricted time of simulation where overlap occurred with varying noise values, for different BK channel conductance values i.e. gBKbar = 150, 300, 450, 600, 750 pS/um2 c) Mutual Information I(R;S) is plotted for restricted time of simulation where overlap occurred with varying noise values, for different high voltage activated calcium channel conductance values i.e. gCaHVAbar= 0.0002, 0.0005, 0.0008, 0.0011, 0.0015 S/cm2 d) Time evolution of mutual information I(R;S) across the whole simulation for different current amplitudes at fixed noise value N=0.2 ‘Colour’ represents current amplitude and ‘line style’ represents type of compartment for the graphs. ‘Blue’ represents 0.4nA current amplitude, ‘red’ represents 0.425nA current amplitude and ‘black’ represents 0.45nA current amplitude. ‘Solid line’ graph represents experimental compartment, ‘dashed line’ represents control 1 and ‘dotted line’ graph represents control 2. e) Time evolution of mutual information for different BK channel conductance values i.e. gBKbar = 150, 300, 450, 600, 750 pS/um2 f) Time evolution of mutual information, for different high voltage activated calcium channel conductance values i.e. gCaHVAbar= 0.0002, 0.0005, 0.0008, 0.0011, 0.0015 S/cm2

**Fig 4:**
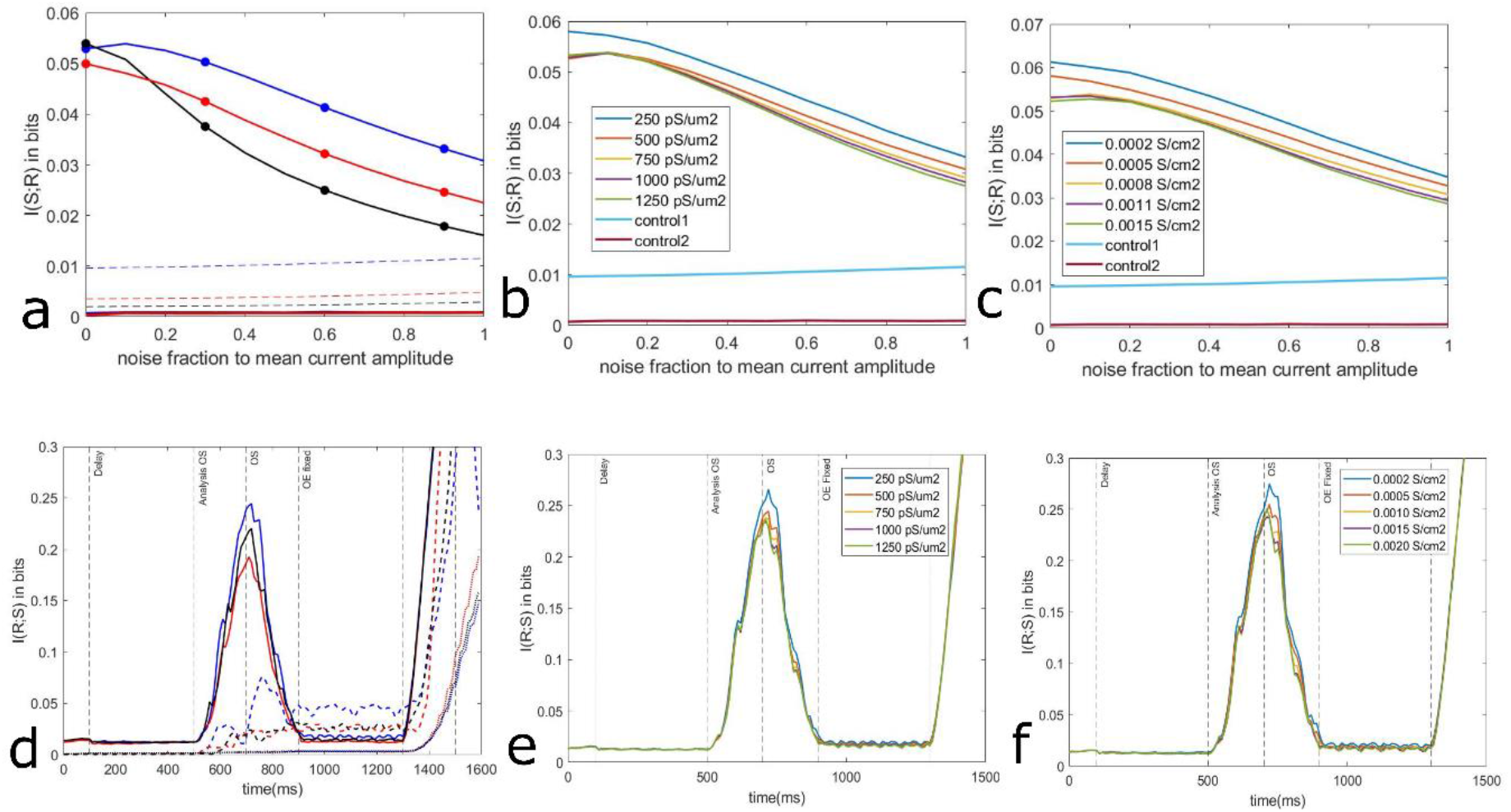
Sensitivity analysis for IPI overlap experiment in dendritic model a) Mutual Information I(R;S) is plotted for restricted time of simulation where overlap occurred with varying noise values, for different current amplitudes of the inputs. ‘Colour’ represents the current amplitude and ‘line style’ represents the nature of compartment i.e. experimental, control 1 or control 2. ‘Blue’ represents current amplitude of 0.l5nA, ‘Black’ represents current amplitude of 0.l75nA, ‘Red’ represents current amplitude of 0.2nA. Graph with ‘filled circle’ markers represent experimental compartment in each colour, ‘dashed line’ represents control 1 and ‘solid line’ represents control 2. Due to less variation in control compartments’ voltage response for different input current amplitudes, the differently coloured graphs are overlapping for both control compartments. b) Mutual Information I(R;S) is plotted for restricted time of simulation where overlap occurred with varying noise values, for different BK channel conductance values i.e. gBKbar = 250, 500, 750, 1000, 1250 pS/um2 c) Mutual Information I(R;S) is plotted for restricted time of simulation where overlap occurred with varying noise values, for different high voltage activated calcium channel conductance values i.e. gCaHVAbar= 0.0002, 0.0005, 0.0010, 0.0015, 0.0020 S/cm2 d) Time evolution of mutual information I(R;S) across the whole simulation for different current amplitudes at fixed noise value N=0.2 ‘Colour’ represents current amplitude and ‘line style’ represents type of compartment for the graphs. ‘Blue’ represents 0.l5nA current amplitude, ‘red’ represents 0.l75nA current amplitude and ‘black’ represents 0.2nA current amplitude. ‘Solid line’ graph represents experimental compartment, ‘dashed line’ represents control 1 and ‘dotted line’ graph represents control 2. e) Time evolution of mutual information for different BK channel conductance values i.e. gBKbar = 250, 500, 750, 1000, 1250 pS/um2 f) Time evolution of mutual information, for different high voltage activated calcium channel conductance values i.e. gCaHVAbar= 0.0002, 0.0005, 0.0010, 0.0015, 0.0020 S/cm2

To make sure that results were robust to variabilities in channel conductance values, sensitivity analysis was done by varying all the inserted channel conductance values for both models. It can be seen from noise varying and time evolving plots for varying BK-channel conductances and Ca-HVA channel conductances, that the information transmission is fairly robust about temporal overlap in the response (Fig 3b, 3e, 3c, 3f for single compartment; Fig 4b, 4e, 4c, 4f). Fast potassium channels, KA channel (dendritic model) and L-type Ca channel conductance values were also varied and the results were same (not shown). Even the sodium and potassium channel conductance values in both controls were varied, and the results remained same (not shown).

### Varying amplitude stimuli input experiment

For this set of experiments in single compartment model, it is seen that even as noise fraction N increases to as high as 3, the mutual information value I(S1,S2;R) is higher (p = 4.988×10^−16 for control1 and p = 3.0795 × 10^−16 for control2) for experimental compartment as compared to both control compartments (Fig 5a). Even for dendritic model similar results were observed for N values as high as 1 (p = 8.9518×10^−4 for control1 and p = 6.6265×10^−4 for control2) (Fig 5c). That means even with as high as 300% noise levels in stimuli amplitudes for single compartment model and 100% noise levels in stimuli amplitudes for dendritic model, the system showing XOR computation optimally transfers information about the two stimuli via voltage response which is very low in the case of control compartments firing constant amplitude APs, both highly excitable and less excitable. It was also observed that the calculation of parameter I(S1,S2;R) provides more insight about the information transmission for two stimuli and one response than that obtained from other measures like I(S1;R) and I(S2;R) in experimental compartment. Whereas the same information measure calculation in control compartment provides very less information about the stimuli and there, the fluctuations are mainly due to noise itself rather than the stimuli (not shown).

**Fig 5:**
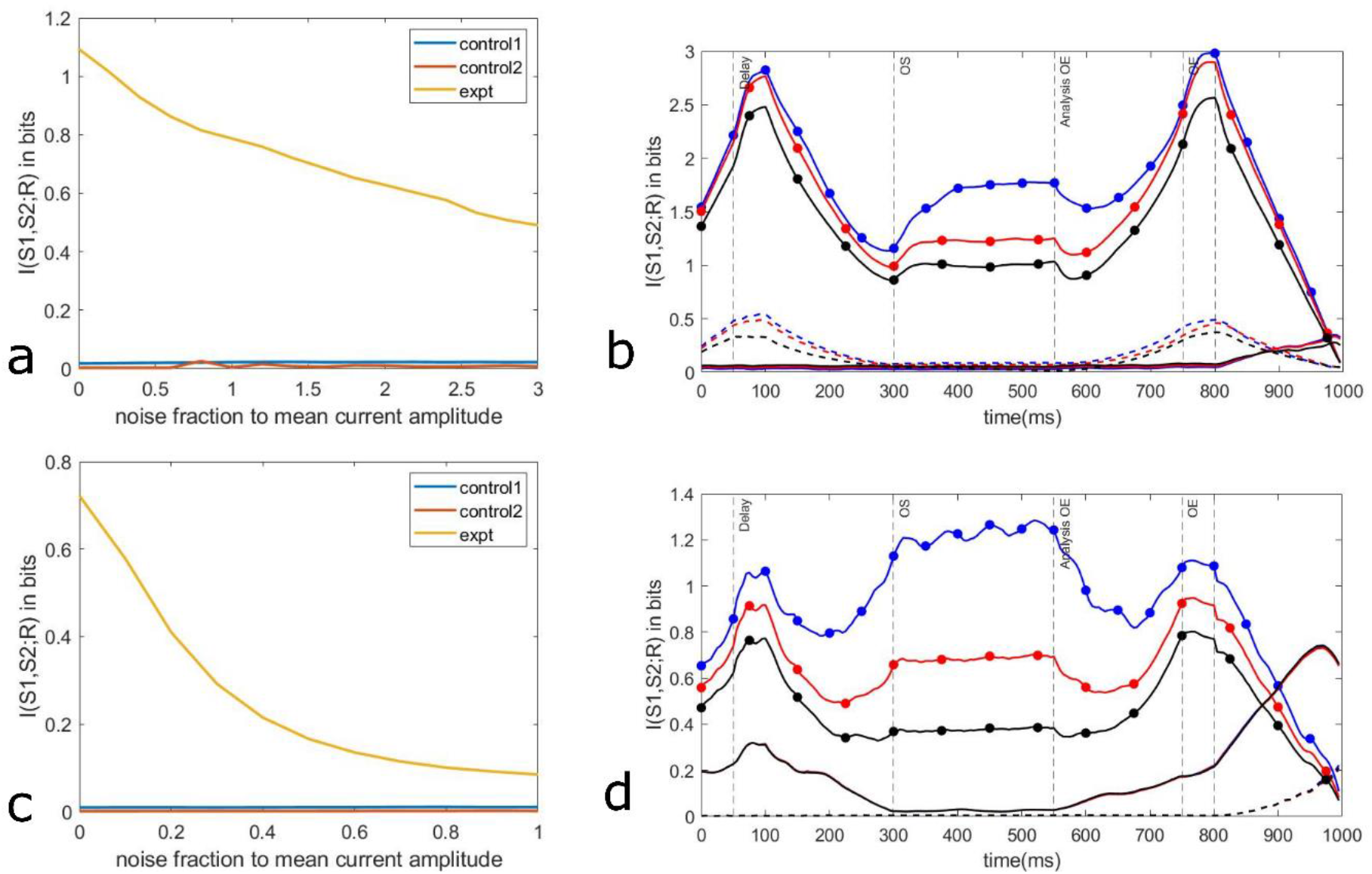
Results from Variable amplitude stimuli input experiment. Mutual Information I(S1, S2; R) is plotted for the whole duration of the simulation with varying noise for single compartment model (a) and dendritic model (c). Time evolution of mutual information I(S1, S2; R) for the whole simulation with different noise values for both single compartment model (b) and dendritic model (d). Colour represents ‘noise value, N’ and ‘line style’ represents nature of compartment i.e. experimental, control 1 or control 2. ‘Blue’ is for N=0.2, ‘Red’ is for N=1 and ‘Black’ is for N=2 in single compartment model whereas, ‘Blue’ is for N=0.1, ‘Red’ is for N=0.3 and ‘Black’ is for N=0.5in dendritic model. Graph with ‘filled circle’ markers represent experimental compartment in each colour, ‘dashed line’ represents control 1 and ‘solid line’ represents control 2.

There is rise of I(S1;R) seen in the duration where stimulus 1 shows in the analysis i.e. from 0 to 300ms whereas I(S2;R) increases during the duration where stimulus 2 shows in the analysis i.e. from 550ms to end of simulation. The increase in these information theoretic parameters increases I(S1,S2;R) (from equation 7), which is the reason we see those peaks during the mention time duration in the graph for I(S1,S2;R) in both models (Fig 5b and Fig 5d). The value of I(S1,S2;R) during the overlap region of time evolution plots i.e. between 300ms and 550ms (even though the actual overlap is from 300ms to 750ms, due to the nature of analysis done in which values for next 200ms are collated, the time at which second stimulus starts showing in I(S1,S2;R) parameter is 550ms) is significantly more than control compartments, and this again reinforces the fact that such a system showing XOR computation is able to transmit the information about multiple variable amplitude input stimuli. It is also seen that as N increases, I(S1,S2;R) value decreases across time which makes sense given the fact that increasing noise does decrease the information about both the stimuli but it is still significantly greater than the control compartments(Fig 5b and Fig 5d).

Sensitivity analysis is done for this set of experiments as well, to make sure the conclusions are robust to changes in parameter values in both models. Similar to IPI overlap experiment, the inserted channel conductance values were varied and all the analysis of experiment repeated. It can be seen from the time evolving and noise varying plots that the most fluctuations were observed when the BK-channel and Ca-HVA channel conductances were varied (Fig 6a, 6b, 6d, 6f in single compartment model; Fig 7a, 7b, 7d, 7f in dendritic model). This is again because these channels are mainly responsible for generating the dCaAPs in both models. But despite that, the conclusions previously drawn remained fairly robust to these variabilities. Noise varying (Fig 6c, 6e in single compartment model; Fig 7c, 7e in dendritic model) and time evolving plots (not shown) for L-type channel, K-fast channel and KA channel (dendritic model) conductances changes showed even less fluctuations.

**Fig 6:**
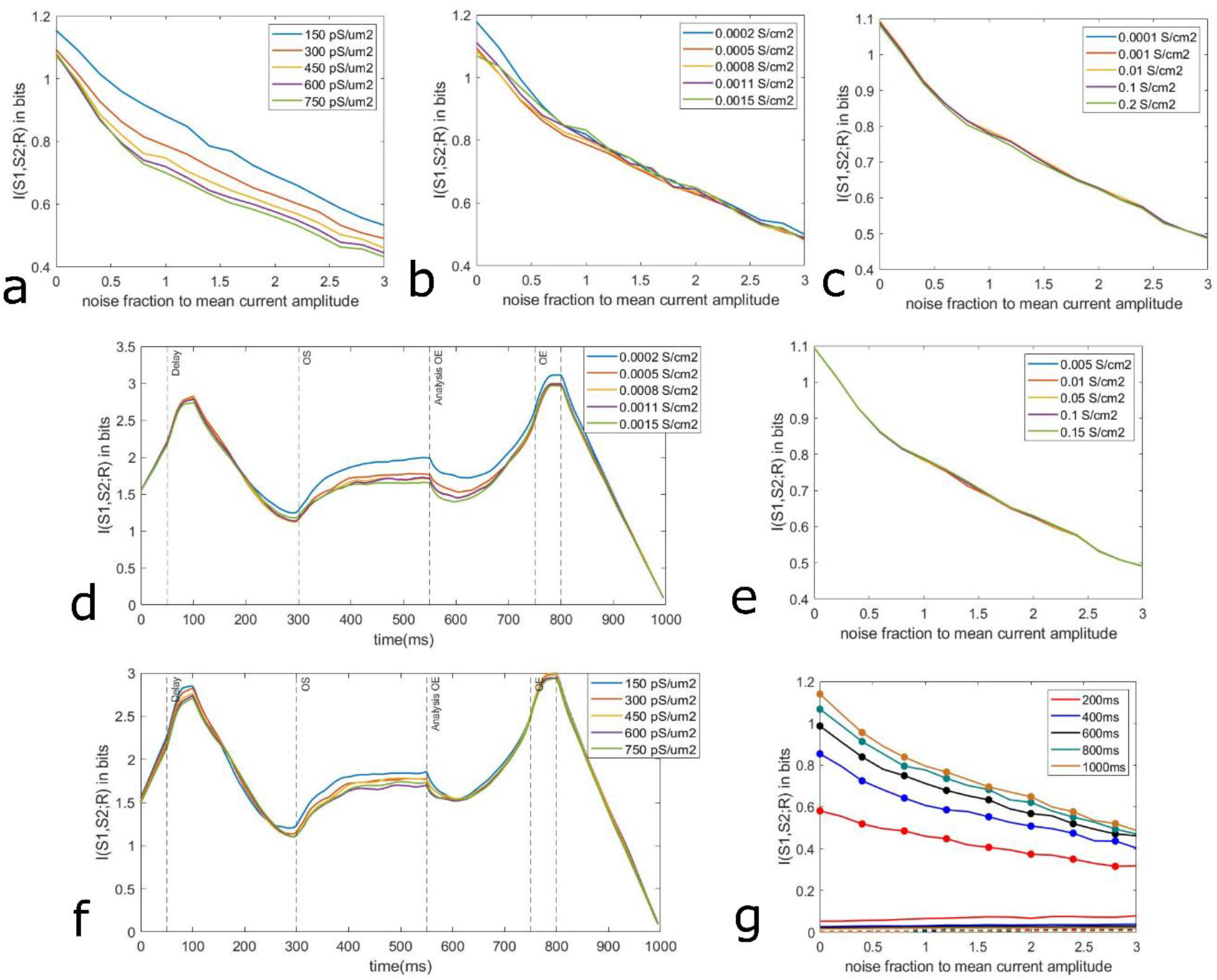
Sensitivity Analysis for Variable Amplitude input experiment in single compartment model a) I(S1,S2;R) is plotted for the whole duration of the simulation with respect to different noise values N on the x axis for different BK channel conductance values i.e. gBKbar = 150, 300, 450, 600, 750 pS/um2 b) I(S1,S2;R) is plotted with N on x-axis for different values of high voltage activated calcium channel conductance i.e. gCaHVAbar= 0.0002, 0.0005, 0.0008, 0.0011, 0.0015 S/cm2 c) I(S1,S2;R) is plotted with N on x-axis for different values of L-type Calcium channel conductance i.e. gCaLbar = 0.0001, 0.001, 0.01, 0.1, 0.2 S/cm2 d) Time evolution of mutual information I(S1, S2; R) for the whole simulation with different values of BK-channel conductance i.e. gBKbar = 150, 300, 450, 600, 750 pS/um2. e) I(S1,S2;R) is plotted with N on x-axis for different values of fast potassium channel conductance i.e. gKfastbar = 0.005, 0.01, 0.05, 0.1, 0.15 S/cm2 f) Time evolution of mutual information I(S1, S2; R) for the whole simulation with different values of high voltage activated calcium channel conductance i.e. gCaHVAbar= 0.0002, 0.0005, 0.0008, 0.0011, 0.0015 S/cm2 g) I(S1,S2;R) is plotted with N on x-axis for different values of temporal overlap i.e. 200, 400, 600, 800, 1000 ms.

**Fig 7:**
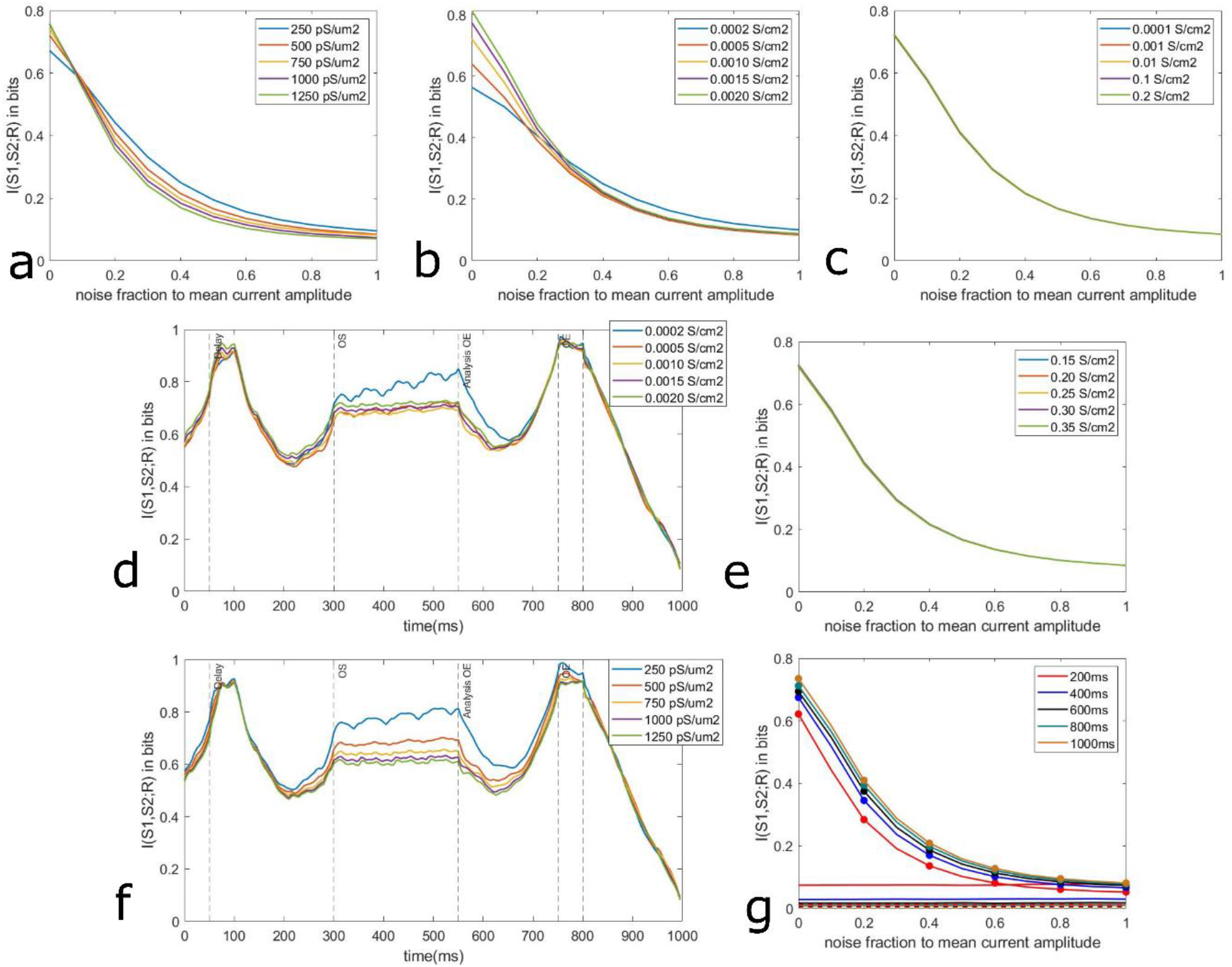
Sensitivity Analysis for Variable Amplitude input experiment in dendritic model a) I(S1,S2;R) is plotted for the whole duration of the simulation with respect to different noise values N on the x axis for different BK channel conductance values i.e. gBKbar = 250, 500, 750, 1000, 1250 pS/um2 b) I(S1,S2;R) is plotted with N on x-axis for different values of high voltage activated calcium channel conductance i.e. gCaHVAbar= 0.0002, 0.0005, 0.0010, 0.0015, 0.0020 S/cm2 c) I(S1,S2;R) is plotted with N on x-axis for different values of L-type Calcium channel conductance i.e. gCaLbar = 0.0001, 0.001, 0.01, 0.1, 0.2 S/cm2 d) Time evolution of mutual information I(S1, S2; R) for the whole simulation with different values of BK-channel conductance i.e. gBKbar = 250, 500, 750, 1000, 1250 pS/um2. e) I(S1,S2;R) is plotted with N on x-axis for different values of fast potassium channel conductance i.e. gKfastbar = 0.15, 0.20, 0.25, 0.30, 0.35 S/cm2 f) Time evolution of mutual information I(S1, S2; R) for the whole simulation with different values of high voltage activated calcium channel conductance i.e. gCaHVAbar= 0.0002, 0.0005, 0.0010, 0.0015, 0.0020 S/cm2 g) I(S1,S2;R) is plotted with N on x-axis for different values of temporal overlap i.e. 200, 400, 600, 800, 1000 ms.

When the temporal overlap window was varied, it was seen from the noise varying plot (Fig 6g single compartment; Fig 7g dendritic model) that as the window decreased, I(S1,S2;R) value across all N values also decreased. This is due to the fact that lesser the temporal overlap window, less time is given for the information transmission about stimuli in the voltage response. For single compartment model and dendritic model, temporal overlap as low as 200ms and 400ms respectively, the mutual information about stimuli in response is greater than control compartments. This again tells us that the results are robust to changes in the value of temporal overlap window parameter.

## Discussion

Early assumptions about existence of only all or none APs have been proved wrong long ago[25]–[28]. Now, even the assumption that dendrites can perform only simple AND, OR logic gates has also been challenged by recent studies showing that XOR computation can be performed on the dendritic level too[9]–[11], [29]. Thus, studying such a system showing graded AP with max amplitude at threshold stimuli and less amplitude for stronger stimuli which can perform XOR computation has provided insights about information transmission about stimuli having different degrees of temporal overlaps as well as stimuli having varying amplitudes. In this study it was observed that even at significantly high noise values, a system showing anti-coincidence phenomenon optimally transfers temporal as well as amplitude information about the stimuli as compared to a control system which doesn’t show this phenomenon.

This fundamental physiological building block can have important implications for sensory system information processing where variable amplitude temporally dynamic noisy inputs are often present[14], [16]. In the dual olfactory pathway of honeybee, projection neurons (PN) transfer information from the primary olfactory neuropile, the antennal lobe (AL) to mushroom body (MB) which is the multimodal integration centre[30]. The PN pathway has mirror-imaged like trajectories i.e., the architecture of two subsystems of AL is counter-rotated with respect to each other[15]. This produces a substantial temporal delay in the downstream neuron of MB[15], [31]. Such a counter-rotating layout is similar to delay lines of vertebrate auditory system, where coincidence detection is used for sound localisation[32]. Thus, it can be proposed that in such systems, XOR computing dendrite can act as a mechanism of information transmission encoding varying degrees of odour concentration and time varying stimuli which are often masked by high degree of noise. Similarly, a possible mechanism for the Kenyon cells to detect different odour concentrations in olfactory system of Drosophila can be via XOR computing neurons since the neuronal architecture is already known to perform coincidence detection[13], [33]. It can also aid in a possible mechanism for stochastic resonance where the precise information about temporal structure i.e., phase of inputs along with amplitude of stimuli is necessary[34].

Such a system may also be able to aid in making the mechanism for BCM synaptic learning more robust[35], [36]. When the pre synaptic inputs are above a certain strength, a dendrite showing this phenomenon would supress the dCaAP amplitude which would ultimately lead to less firing of neuron i.e., less feedback from soma to the particular synapse. This will lead to lowering of synaptic modification threshold. When the pre-synaptic input would be lesser than the strength (above which XOR phenomenon is seen), increasing pre-synaptic input amplitude would lead to increased post synaptic response which means more firing of neuron in that range i.e., more feedback from soma to that synapse leading to increase in synaptic modification threshold. This dendritic tuft XOR computation along with AND/OR computations at soma level in cortical neurons can then together act a strong fundamental computational block of a single neuron affecting plasticity rules in a wide range of possibilities[37]–[39]. In connection with the recent experimental finding on such systems[9], such information theoretic computational approaches[4], [13], [40] may help in finding out the role of new phenomenon and what kind of conditions govern optimal information transmission.

